# Continuous Temporal Difference Learning as a Unifying Theory of Dopamine Function

**DOI:** 10.64898/2026.04.07.716149

**Authors:** Sankalp Garud, Laurel Morris

## Abstract

Dopamine neurons are thought to signal reward prediction errors phasically and the opportunity cost of time tonically, while also displaying ramping activity during goal approach, and coupling with movement. These are often treated as distinct modes of dopamine function, each requiring its own computational explanation. Here we show that all can be unified by considering temporal difference learning within the context of continuous time, combined with the assumption that the brain computes value changes through a fast model-based process while maintaining a slower model-free cache. Together, the inclusion of these two ingredients explains phasic responses, tonic modulation between reward contexts, navigation ramps, speed scaling, and the fading of ramps with learning, without invoking separate mechanisms. We confirm these predictions across two independent datasets of dopamine recordings in rodents spanning freely-moving and head-fixed paradigms. Continuous temporal difference learning may thus provide a unified theory of dopamine function.

## INTRODUCTION

Dopamine neurons in the mammalian midbrain display a diversity of signalling patterns. They fire phasic bursts at unexpected rewards and cues that predict rewards^1^, ramp up their activity as an animal approaches its goals^2,3^, represent reward contexts in their tonic baselines^4–7^, and even co-vary with movement, speed, or force when interacting with valued outcomes^8–10^. Each have been treated as different, often contradictory, modes of dopamine function, and together they have motivated a longstanding debate about the fundamental computational basis of dopamine function^11^.

The most successful account of these signals has come from temporal difference (TD) learning, which proposes that dopamine neurons signal reward prediction errors, the difference between received and expected reward^1^. This framework explains a remarkable range of phasic dopamine responses: bursts to unexpected rewards, transfer of responses from reward to reward-predicting cues, and dips at reward omission. Some recent findings, however, challenge the RPE account. Dopamine in the ventral striatum ramps up over seconds as animals approach reward^2^, with the ramp scaling with reward magnitude and proximity^3^. This has led some to suggest that dopamine might not represent prediction errors at all, but something different, such as value^12^, force^10^, or velocity^13,14^. Separately, tonic dopamine, modulated by reward context and linked to behavioural vigour, has been treated as its own signalling mode with its own computational explanation^4^.

Most models of dopamine function are inspired by reinforcement learning algorithms, where artificial agents learn by stepping between discrete states and receiving rewards at transitions^15,16^. This discretisation is required for artificial systems that train over epochs, but animals experience continuous time, a continuous stream of sensory information, and a continuum of states. Whether an animal runs or walks between two locations, the prediction error in a discrete model is the same. We suggest that taking continuous states and time into account may be important, and similar to other works^3,17^, dopamine ramps might reflect a derivative-like computation.

Here, we propose a unifying model of dopamine function by considering the continuous-time formulation of temporal difference learning, applying the chain rule, and incorporating the well-motivated distinction between model-based and model-free value estimates, a key finding of recent computational and experimental work^18–20^. Our framework accounts for ramps during goal approach, scaling with movement speed, tonic modulation across reward contexts, and the fading of ramps with learning. It also generates novel predictions, for example, that dopamine should dip below baseline during stationary periods. We test these against two independent datasets spanning freely-moving and head-fixed paradigms.

## RESULTS

We first start by outlining the continuous time temporal difference (TD) learning theory, and contrasting it with its discrete counterpart. First, in discrete time, the reward prediction error is given by:

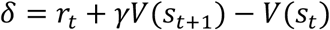

In continuous time, this becomes^21^:

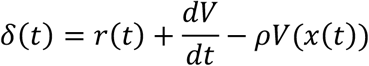

where *V*(*x*) is the value function contingent on a time dependent state x, *r*(*t*) is the reward function, *ρ* is the rate of discount (see Supplementary S1 for a full derivation of the framework).

Applying the chain rule, we get:

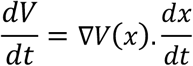

In the case of a rat approaching a goal, ∇*V*(*x*) could be interpreted as a spatial gradient which varies with its position x(t). Similarly, 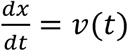, would denote the animal’s instantaneous velocity.

Crucially, the terms in the continuous temporal difference error equation need not share the same computational origin or architecture^22^. In line with recent studies showing striatal dopamine representing both model based and model free estimates^18,20,23^, we propose that the spatial gradient ∇*V*(*x*) could be derived from a model-based map of the environment, available as soon as the animal learns the task structure. On the other hand, we propose that the value estimate 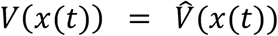 reflects a cached model-free representation, updated incrementally through direct experience.

Therefore, the continuous time temporal difference error can be written as:

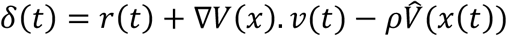

When an animal approaches a reward, discrete TD models would predict a sharp phasic spike at the point of reward delivery, which would propagate back to subsequent states with learning (Fig 1B). In continuous time, the prediction error becomes a smooth continuous signal, resembling a ramp (Fig 1C). The shape of the ramp arises from the gradient term ∇*V*. For a single reward at a fixed location, TD learning with, say, exponential discounting, produces a value function that rises toward the goal. Consequently, its spatial derivative would steepen in proximity to the goal. Multiplied by the animal’s velocity, this generates the characteristic ramp. This is consistent with, and provides a precise account of, previous suggestions proposing that striatal dopamine performs a derivative like calculation^3,17^.

**Figure 1.**
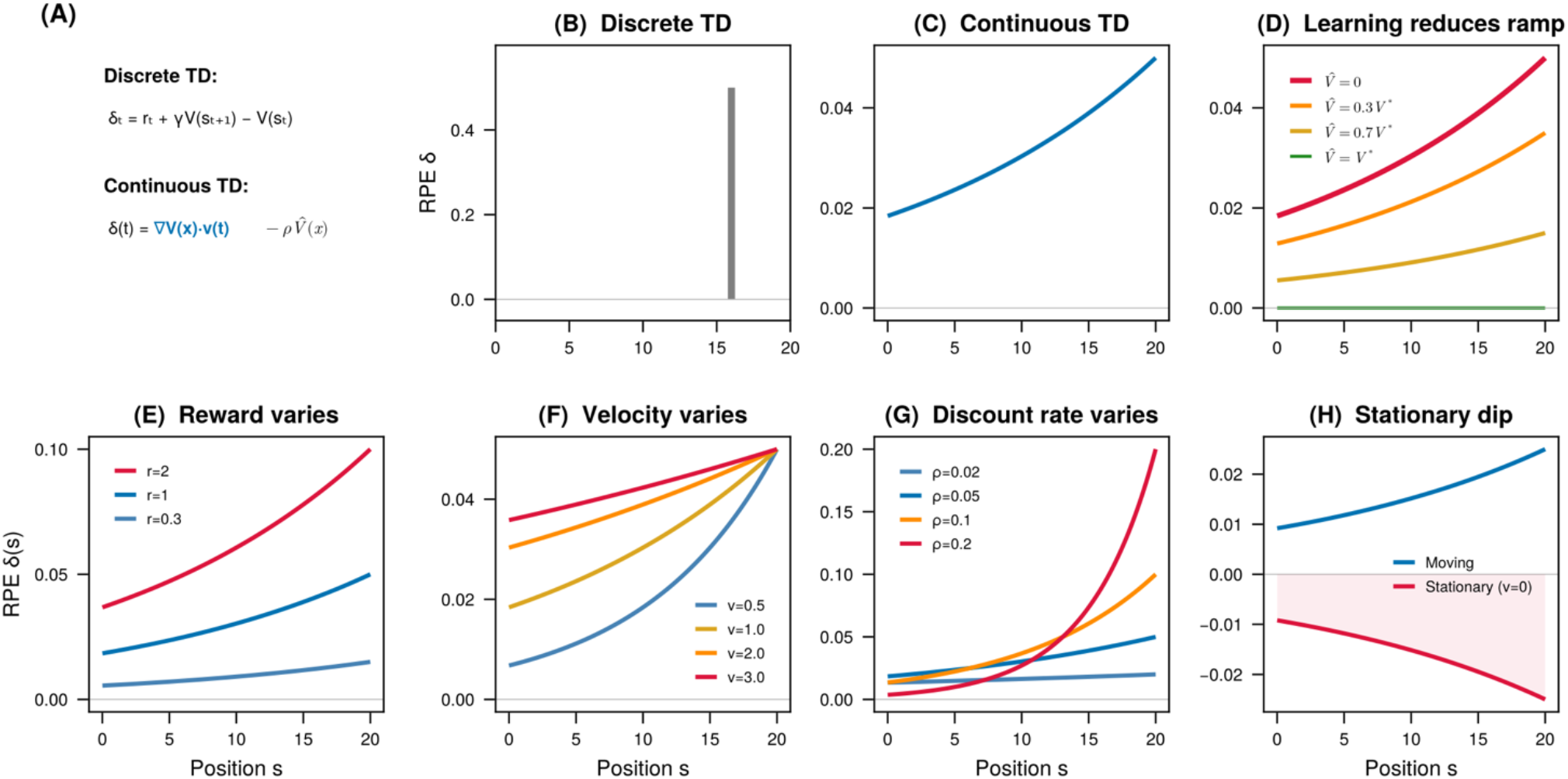
The continuous time TD model and its predictions. (A) The discrete and continuous TD-error, with the blue text indicating a chain rule transformation that would produce a ramp. (B) Predictions from a discrete time RPE model. In a discrete TD model, the RPE is a point prediction error that propagates backward one state per update. (C) In continuous TD, the chain rule produces a spatial ramp, assuming the value function is sufficiently convex. (D) As learned value converges to true value, the model predicts that the ramp would flatten with learning. E) The model predicts that the ramp will scale with reward magnitude, (F) velocity, and (G) the temporal discount rate. (H) The model predicts a dip when the animal is at rest, as the velocity is zero and the negative term dominates.

As the cached value 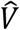 converges to the true value, the ramp begins to flatten (Fig 1D). Similarly, ramp scales with increasing reward, velocity, and steepens with higher discount rate (Fig 1E-G). Importantly, our model predicts that at rest, the temporal difference error dips below baseline, and the negative term −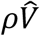dominates (Fig 1H).

We tested these predictions against the data from Krausz et al (2023)^24^, who recorded nucleus accumbens dLight photometry in freely moving rats approaching a reward port. As noted in many past studies^2,3,24^, and consistent with our model predictions (Fig. 1C), DA ramped as animals approached their goals (Fig. 2A). Moreover, the ramps scaled with reward magnitudes, and velocity (Fig. 2B, D)., as predicted by our model (Fig. 1E, F) 1)In this study, the rats faced complex environmental contingencies, meaning learning did not get to converge, and therefore the ramps in early and late sessions look similar (i.e. the ramps do not flatten across time in the absence of learning). We found that the velocity co-varied with the distance to goal (Fig. 2E). Interestingly, the velocity-dopamine coupling increased with proximity to goal, consistent with the idea that velocity scales with the gradient (Fig 2F, r=-0.494, df=18, p = 0.027).

**Figure 2.**
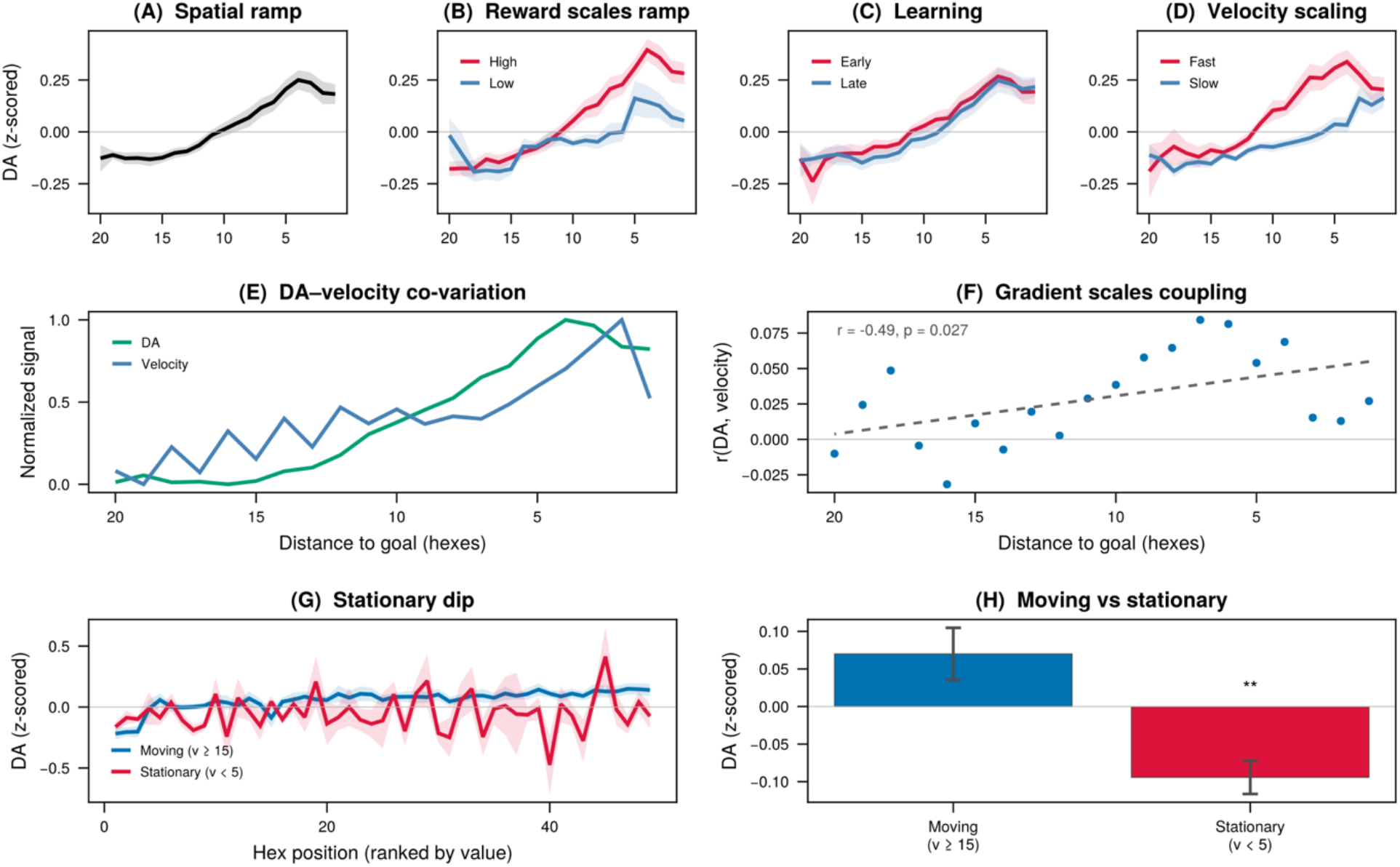
Predictions of continuous time TD error hold in goal seeking rats. (A) DA ramps as the rats approach a reward port, consistent with the value gradient (B) The ramp is steeper on approaches ending at high reward (>80 µL juice) versus low reward (< 20 µL), consistent with the spatial gradient scaling with the reward magnitude. (C) Per-rat quartile split on session number shows a modest reduction in ramp amplitude from early to late sessions. (D) Per-rat quartile split on velocity: fast trials show larger DA at the same positions, consistent with velocity scaling the ramp. E) Average dopamine and velocity covary across the distance to goal. (F) Velocity and DA are linked stronger toward the goal, consistent with the idea of a spatial gradient modulating this relationship. (G-H) Stationary DA dips below zero as predicted when considering both approach and non-approach trials. All panels use NAc dLight photometry data from Krausz et. al. 2023.

One prediction our model uniquely makes is the DA values would dip below zero at points where the rat’s velocity tended to zero (being stationary). We separated the DA signals into stationary (v <5 cm/s) and rapidly moving instances (v >15 cm/s), and found that indeed the dopamine dipped below baseline (df =9, t = -4.27, p = 0.001, one-tailed) in instances where the rats’ velocity was near zero (Fig 2G-H).

A key algebraic property of the model affords us the opportunity to test its internal consistency. When an animal is moving, DA reflects 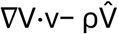; when stationary (v = 0), DA reflects only −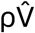. Subtracting stationary DA from moving DA at matched positions cancels the −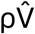 term, isolating ∇V·v. Dividing by velocity yields the spatial gradient ∇V, which can then be numerically integrated to recover V(x) without free parameters (Fig 3A-B).

**Fig 3.**
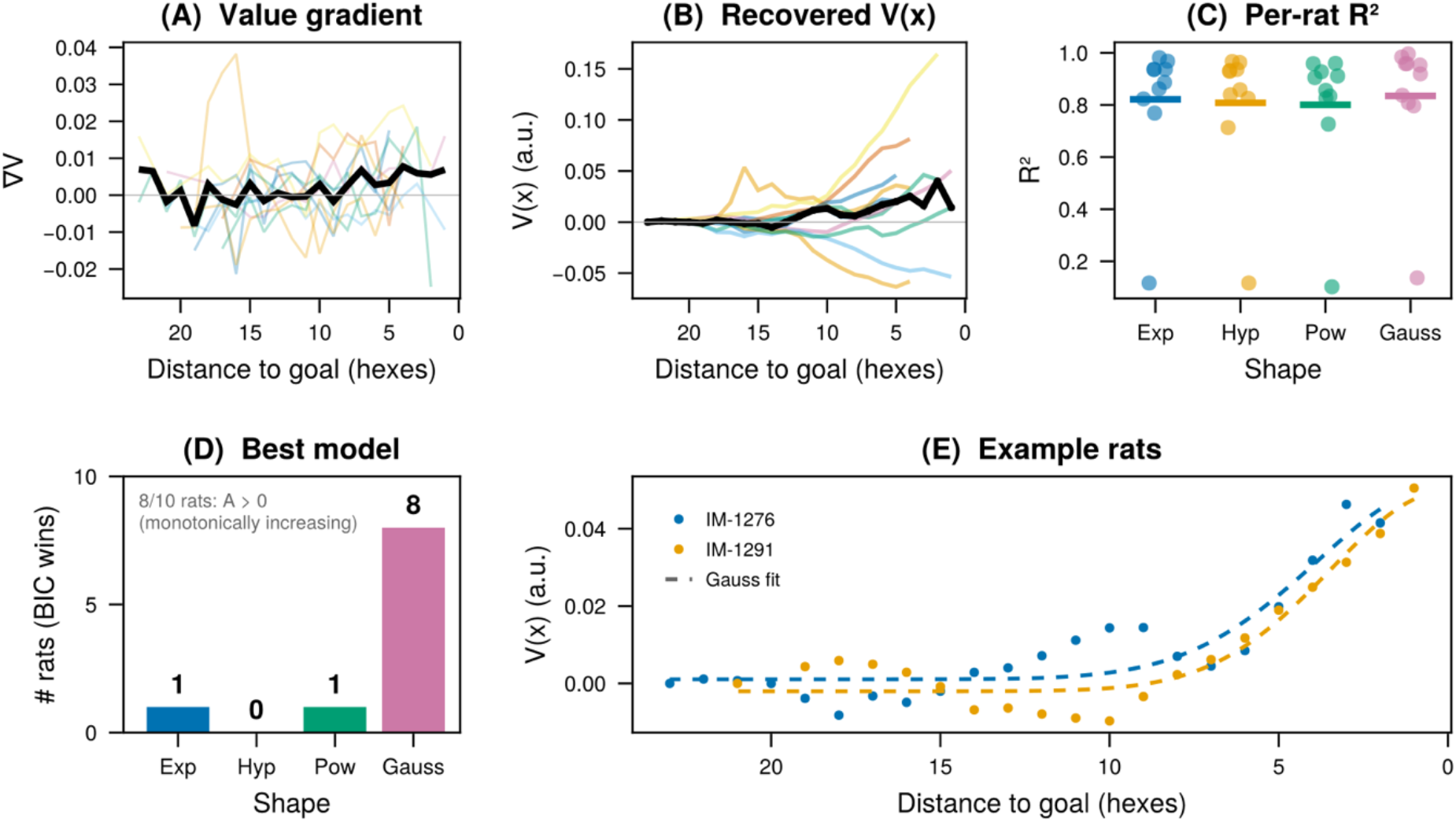
Internal Consistency: Recovering V(x) from the dopamine signal suggests it increases monotonically. (A) Panel showing the value gradient (B) The recovered value function increases in proximity to the goal. (B) Four models are fitted to the value function, but a gaussian model fits the best (C-D) in most rats. (E) Example rats’ value functions fitted with the exponential and gauss TD model.

The recovered value function showed a net increase toward the goal in 8 out of 10 rats (A>0 in BIC winning model). We fitted four candidate shapes — exponential, hyperbolic, power-law, and Gaussian — to each rat’s recovered V(x). All four provided good fits (mean R^2^ > 0.8), with the best-fitting shape varying across rats: a Gaussian profile won in 8, exponential in 1, and power-law in 1 (Fig 3C-D). This individual variability suggests that the value function might be monotonically increasing, suggesting individual differences in how spatial value is represented.

We next focus our attention on the negative term (−*ρV*), and show how our model might offer a concrete interpretation of the tonic dopamine signals. In our model, when no reward is being delivered and the state variable is not changing, the gradient and reward terms vanish, leaving only the negative cached value scaled by the discount rate. Crucially, the state variable need not be absent; the animal may well have learned the value of its current state. Under these conditions, the residual dopamine signal is what might be considered tonic dopamine, and our model predicts that it reflects the negative cached value. If so, this tonic signal should deepen as the animal learns, since the cached value grows with experience, and behavioural vigour should increase in parallel, following proposals that tonic dopamine might drive behavioural vigour^4^. To examine this, we used dLight photometry recordings from head-fixed mice performing a Pavlovian conditioning task with short (∼8s) or long (∼55s) inter-trial intervals (Floeder et al., 2025; DANDI:001552), focusing on the late inter-trial interval, after reward has been consumed and before the next cue appears.

Mice in short-ITI sessions showed higher anticipatory lick rates (Fig. 4A), consistent with Niv’s (2007) prediction that the average reward rate sets behavioural vigour. During the ITI itself, long-ITI dopamine settled reliably below baseline — the tonic dip predicted by −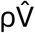— while the short-ITI trace showed a slowly decaying reward response that did not resolve before the next trial (Fig. 4B). Focusing on the long-ITI condition where the tonic floor is cleanly measurable, we found that the dip deepened across training sessions as 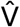was learned (Fig. 4D, r=-0.29, p<1e-3), while anticipatory vigour increased in parallel (Fig. 4C, r = 0.65, p=so this 1e-4). These results suggest that tonic dopamine might encode an ongoing opportunity cost signal −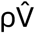that deepens with learning and covaries with behavioural vigour.

**Fig 4.**
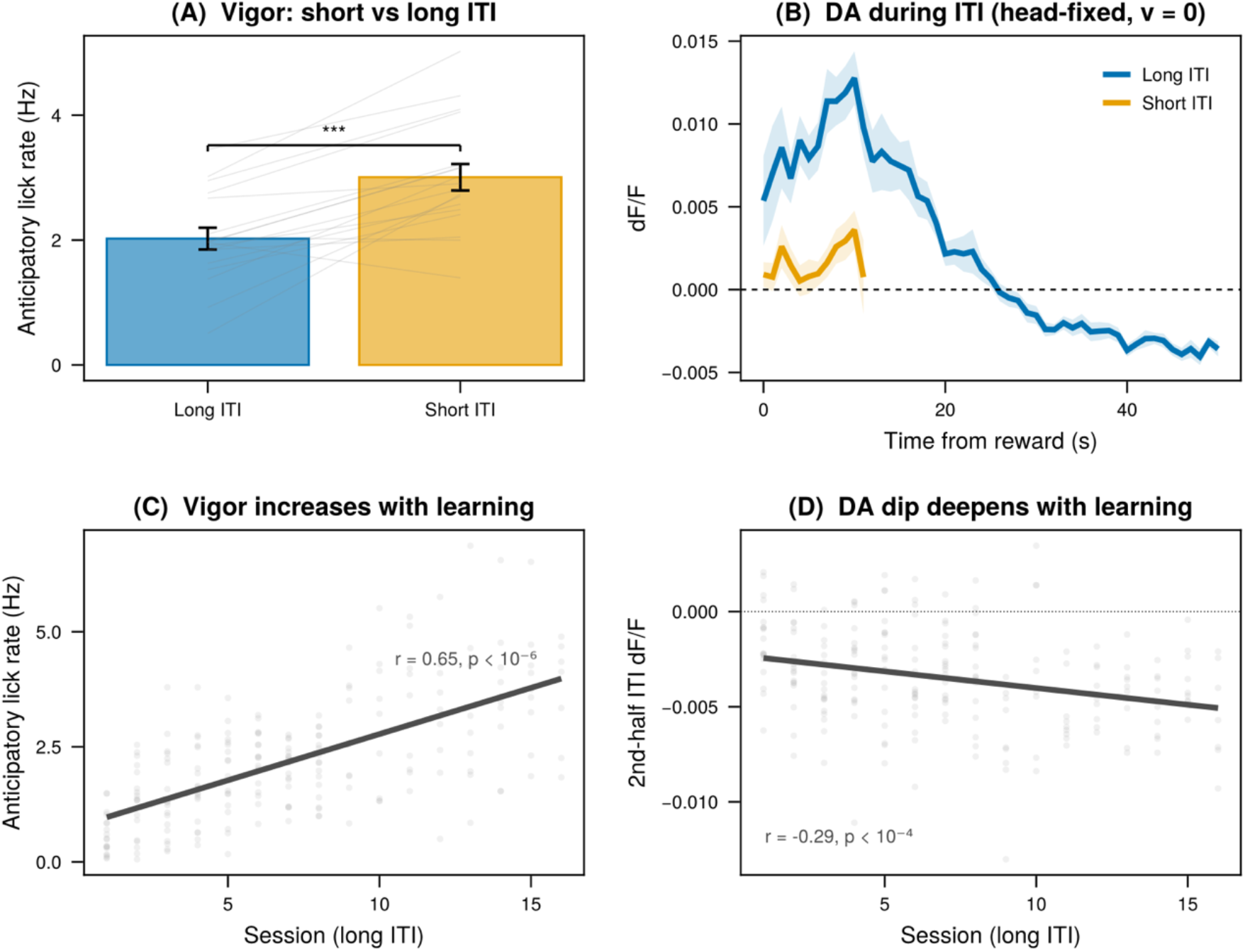
Continuous RPE model explains vigour and tonic DA function. (A) Anticipatory lick rate is higher in shorter compared to longer ITI sessions. (B) DA time course following reward delivery. Long ITI sessions reveal a dip below baseline, revealing a tonic signal consistent with the model prediction. (C) Lick rate increases with long ITI sessions, indicating growing vigour with training. (D-E) Conversely, DA dips deepen with learning, consistent with the model prediction, and with theories of tonic dopamine as reflecting the opportunity cost of time. All panels use dLight photometry from Floeder et al. (2025).

Finally, we compared the continuous-time TD RPE model quantitatively against competing accounts of dopamine during approach. We fit six models to per-rat, observation-level data: two variants of our CT TD model (exponential and Gaussian spatial profiles, both including the tonic 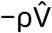 term), two corresponding dV/dt models that omit the tonic term, a pure value model, and a velocity-only model. Fig. 5A illustrates the qualitative predictions of each model class on a simulated approach-with-pause trajectory. Only the CT TD model predicts both a velocity-scaled ramp during movement and a negative deflection during the pause.

**Fig 5.**
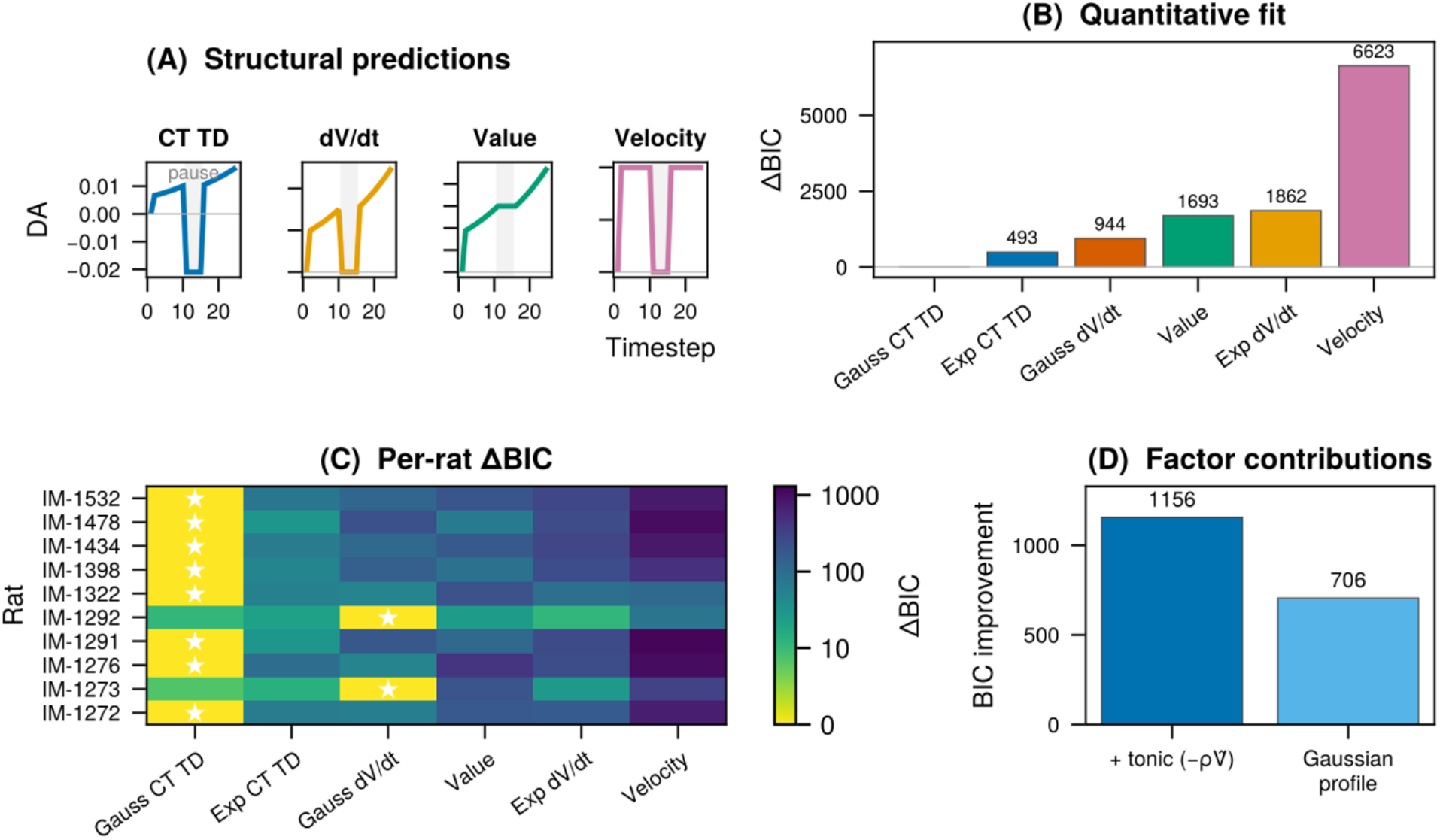
Quantitative model comparison across six competing accounts. (A) Structural predictions for an approach-then-pause trajectory; only CT TD predicts a dip below baseline during the pause. (B) Summed ΔBIC across rats favours the Gaussian CT TD model. (C) Per-rat ΔBIC heatmap confirms consistency across individual animals. (D) Both the tonic term and Gaussian spatial profile contribute independently to model performance.

The Gaussian CT TD model provided the best fit across rats (Fig. 5B, Krausz 2023 dataset), and was also winning in 8 of 10 individual rats by BIC (Fig. 5C). To illustrate the contribution of each component, we decomposed the four CT TD / dV/dt variants factorially (Fig. 5D). The tonic term −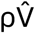improved fit by ΔBIC =1156, while the Gaussian profile improved fit by ΔBIC = 706. Both contribute substantially, but the tonic term (which captures the stationary dip below baseline) accounts for the larger share.

Taken together, these results suggest that a single model can account for dopamine signals that have previously been attributed to distinct computational mechanisms. The chain rule decomposition separates the RPE into three terms, each of which can dominate under different behavioural conditions: during goal approach, ∇V · v may produce ramps that scale with the rate of state change; during rest periods, this term vanishes and the tonic cost − 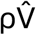is exposed; at reward delivery, r(t) produces the classical phasic burst. Table 1 formalises these predictions across five behavioural regimes. Rather than requiring separate theories for ramping, velocity coupling, tonic modulation, and phasic responses, each may emerge as a different operating point of the same computation, governed by which term dominates under the prevailing conditions. Indeed, apart from the findings described in the main text, the framework continues to describe the classical phasic conditioning response (see Supplementary Figure S1 for a walkthrough), and correctly predicts no dopamine-velocity coupling in the absence of a spatial value gradient (confirmed in an independent dataset of mice running without a spatial goal^25^, Supplementary Figure S2).

**Table 1.** One equation, multiple regimes. The continuous-time contains three terms — instantaneous reward, the value gradient scaled by velocity, and the opportunity cost of the current state — whose relative magnitudes determine the predicted DA signal in each behavioural regime.

| Condition | $r(t)$ | $\nabla V \cdot v$ | $-\rho \hat{V}$ | Predicted DA |
| --- | --- | --- | --- | --- |
| Approaching goal (moving) | 0 | Dominant | Small | Ramp |
| Paused near goal | 0 | Zero | Dominant | Dip |
| Running (no spatial goal) | 0 | $\propto v(t)$ | Small | Tracks speed |
| Stationary (rest / ITI) | 0 | Zero | Dominant | Suppressed |
| Reward delivery | Dominant | Varies | Small | Burst |

Blue: $\nabla V \cdot v$ dominates. Red: $-\rho \hat{V}$ dominates. Green: $r(t)$ dominates. Setting $v = 0$ switches between the first two.

## DISCUSSION

In this study, we showed that by considering temporal difference learning in continuous time^21^, and incorporating the distinction between model-based and model-free value estimates, a single model might account for what have previously been treated as separate dopamine functions. The continuous-time formulation yields a prediction error with three terms — reward, the rate of change of value, and a discounted cached value — and applying the chain rule to the rate-of-change term introduces velocity and a spatial gradient directly into the prediction error. Together, these terms explain dopamine ramps during goal approach, their scaling with speed, a dip below baseline during stationary periods, tonic modulation across reward contexts, and the fading of ramps with learning. We confirmed these predictions across two independent datasets spanning freely moving and head-fixed paradigms.

Consistent with previous studies^18–20,23^, our model proposes that the dopamine signal reflects both model-based and model-free contributions, and that a chain rule-like computation in the brain may make the former tractable. The chain rule is not merely an algebraic rewrite; it is a computational decomposition that expresses the rate of change of a function in terms of other functions, which for a rat navigating toward a goal might mean the computation of a spatial gradient and a velocity term. The chain rule may enable a decomposition of abstract value in terms of quantities already available to the brain — position, if the brain maintains a map of the environment, and velocity, from motor and proprioceptive systems. The decomposition thus reframes a simulation problem as a retrieval problem: the relevant quantities are already represented, and therefore computable, without requiring the brain to mentally roll out future states.

In the case of a rat approaching a reward, the chain rule decomposes the rate of change of value into a spatial gradient and velocity. This accounts for three features of dopamine during goal approach (Fig. 2): the ramp itself (a positive gradient multiplied by a positive velocity), its scaling with speed (faster movement amplifies the signal), and the dip below baseline when the animal is stationary (the velocity term vanishes, exposing only the negative cached value term). We confirmed these predictions in the Krausz et al. (2023) dataset.

Our model would predict that ramps are largest early in learning and diminish with training. This prediction follows from the dual-system assumption: the spatial gradient, derived from model-based maps, updates once the animal learns the task structure. On the other hand, the cached value estimate is maintained through trial-and-error learning, and lags behind model-based estimates. The difference between model based and model free estimates produce the ramp, which eventually attenuates as the model free term converges to true value. This is consistent with empirical work showing that dopamine ramps fade with learning^3^, and a recent theoretical work that shows dual-process architecture explains dopamine ramps which fade with learning, and re-emerge after environmental contingencies change^23^.

Our framework affords an internal consistency check (Fig. 3). When the animal is moving, the dopamine signal reflects the value gradient scaled by velocity minus the cached value. When the animal is stationary at the same position, only the negative cached value remains. Subtracting stationary dopamine from moving dopamine at matched positions cancels the cached value term exactly, isolating the gradient scaled by velocity. Dividing by velocity yields the gradient itself, which can be integrated to recover the value function without fitting any free parameters.

We found that the recovered value function is consistent with monotonically increasing functions, thus suggesting our framework is internally consistent. The best-fitting spatial profile across animals is smooth and Gaussian-like, rather than strictly exponential. Importantly, the recovery procedure is agnostic to the shape of the value function: it does not assume exponential discounting or any other parametric form, but recovers whatever spatial profile the brain maintains. More broadly, the framework is agnostic to the representation underlying the value function — whether computed via exponential discounting, successor representations, or belief-state maps. That the recovered shape is smooth rather than sharply exponential may be consistent with belief-based value function proposals^20^, in which value is computed not over exact position but over the animal’s uncertain estimate of where it is. Under such a representation, spatial uncertainty would naturally smooth the value function relative to the sharp exponential predicted by a model that assumes complete knowledge of position. The framework does not depend on this interpretation, but the recovered shape is consistent with what one would expect if the brain represents value over beliefs rather than states.

During periods where state variables are unlikely to be progressing, for example, late intertrial intervals in head-fixed paradigms (Fig. 4), the gradient and reward terms might be absent leaving only the negative cached value. This provides a clean window onto what has traditionally been called the tonic dopamine signal, isolated from the ramp-generating gradient term. Using photometry recordings from head-fixed mice performing Pavlovian conditioning with short and long inter-trial intervals, we find that dopamine during the inter-trial interval settles reliably below baseline, the negative deflection predicted by the cached value term. This dip deepens across training sessions as the cached value is learned, while anticipatory licking increases in parallel, consistent with the proposal that tonic dopamine reflects the opportunity cost of inaction^4^.

This result suggests that tonic and phasic dopamine may not be separate signalling modes requiring distinct computational explanations. Instead, they may represent the same prediction error observed at different operating points: when the animal is moving, the gradient term dominates and produces ramps and phasic-like transients; when the animal is stationary, the gradient term vanishes and the cached value term is exposed as a tonic baseline shift. The longstanding distinction between tonic and phasic dopamine function may thus reflect not two systems but distinct regimes of one computation (Table 1).

The continuous-time temporal difference framework outperforms derivative-only, value-only, and velocity-only models in a quantitative comparison across individual rats (Fig. 5). The cached value term carries substantial explanatory weight, indicating that it is not a negligible correction but a major component of the dopamine signal.

Previous work correctly identified that dopamine during navigation performs a derivate-like computation^3,17^. By considering the continuous form of temporal difference error, our framework shows why a derivative computation might be necessary in principle if indeed dopamine reflected temporal difference errors. The chain rule decomposition goes further by showing separating the spatial gradient from velocity and exposing the cached value term, neither of which is visible in the undifferentiated derivative. This separation matters empirically: velocity and the gradient make distinct, testable contributions, and the cached value term explains the stationary dip and tonic modulation that a pure derivative account does not predict.

An opposing proposal has been that dopamine reflects value signals, not prediction errors^12^. In our framework, value coding is what the gradient-times-velocity term looks like when speed is roughly constant. Under constant velocity, the prediction error is approximately proportional to the value gradient, which in turn tracks value. Value coding is therefore not inaccurate but perhaps incomplete: it captures the signal under restricted conditions, and the quantitative comparison confirms that including velocity appreciably improves fit.

A seemingly opposite line of work suggests that dopamine signals force during instrumental behaviour, not prediction errors^10^. Since the derivate of velocity is acceleration, our framework naturally predicts a relationship between dopamine and force (force is mass times acceleration according to Newton’s second law of motion). Importantly, force coding need not be a separate framework; it may arise naturally from considering a continuous-time TD model, as the gradient-times-velocity term will covary with any quantity that covaries with velocity, including force.

To summarise the predictions of the framework in different behavioural regimes: the continuous-time prediction error contains three terms, reward, the value gradient scaled by velocity, and the cached value, whose relative magnitudes determine the predicted dopamine signal in each regime (Table 1). During goal approach, the gradient term dominates and produces ramps. When the animal is stationary, the gradient vanishes and the negative cached value is exposed. At reward delivery, the reward term produces the classical phasic burst. Between reward contexts, the cached value reflects the average reward rate. And in the absence of a spatial goal, the gradient is zero and dopamine should not track velocity at all, a prediction we confirmed in a third independent dataset of freely running mice with no position-dependent reward contingency (reported in supplementary analysis)^25^. Each of these regimes has previously been treated as requiring a separate computational explanation. The present work suggests they may instead represent different operating points of one computation, governed by which term dominates under the prevailing behavioural conditions.

We note two principal limitations. First, we do not test trial-by-trial learning trajectories. The framework predicts that ramps should attenuate as the cached value converges toward the true value, and this is consistent with published reports of ramp reduction with overtraining, but we have not fitted the convergence dynamics directly. The Krausz dataset involves dynamic reward contingencies that make convergence tests difficult. Second, the dual-system assumption, that the gradient and cached value are maintained by separate learning systems with different update rates, is theoretically motivated and consistent with the broader model-based/model-free literature, but future works need to empirically verify its bases in the dopaminergic midbrain.

One of the prominent findings of the recent past has been the heterogeneity of dopamine neurons^26^. One way to test the dual-system assumption empirically might be to examine the structure of this heterogeneity. If the prediction error is assembled from factorised components, then dopamine neurons should be heterogeneous in a structured way. For instance, neurons receiving stronger hippocampal input should be more gradient-like, modulated by position and sensitive to reward location. Neurons receiving stronger motor or proprioceptive input should track velocity. Neurons reflecting the cached value should be the slowest to update across learning and the most dependent on direct experience. In this view, heterogeneity is not noise or redundancy but may be a computational requirement: computing a chain rule over terms that arrive from different brain regions on different timescales may demand a population with structured input diversity. Multi-unit recording combined with input tracing techniques could test these predictions directly.

The dual system assumption could also be tested causally. Inactivating or adding noise to hippocampal or prefrontal inputs to the dopamine system should selectively alter the model-based gradient component, eliminate ramps and velocity scaling, while leaving the cached value term intact, preserving phasic responses to reward and the tonic baseline shift.

Crucially, our framework makes no assumption about what the state variable is. In freely moving tasks, the relevant state is physical position and the velocity is locomotor speed. In head-fixed preparations with dynamic sensory cues, the state could be sensory or cue space, with velocity reflecting the rate of sensory change. The chain rule applies wherever the brain maintains a value function over some progressing state variable, and this flexibility means the framework is not tied to spatial navigation but extends to any domain in which value changes over time.

The foundational algorithms that have shaped artificial intelligence, including reinforcement learning, were designed for software agents stepping between discrete states, choosing actions at each state, and receiving rewards at transitions. This discretisation was necessary for tractable value updating, learning, and credit assignment in artificial systems. Such algorithms have been extraordinarily productive in creating artificial agents and explaining biological ones alike, but they carry an implicit assumption: that time is experienced as a sequence of discrete moments, and that the prediction error is a punctate event occurring at each step. Animals, however, do not experience the world in discrete states. Instead, they move through continuous space, receive a continuous stream of sensory information, and act in continuous time. In this study, we have suggested that by considering continuous reinforcement learning, apparently disparate functions of dopamine may be understood as one signal, observed under different conditions.

## METHODS

### Datasets

Two datasets (with a third dataset used in the supplements) were used to analyse predictions from this model. In the first dataset used to characterise ramping DA signals, Krausz et al.^24,27^ recorded dopamine activity in the NAc using dLight photometry. The rats (n=10) navigated a hexagonal grid, which comprised 49 hex positions in a triangular structure. There were three reward ports at the vertices of the triangle, and the rewards magnitudes varied across blocks. Each rat completed 5-12 sessions (82 rat-sessions total). The dataset provided hex-level observations of z-scored DA, instantaneous velocity, distance to chosen reward port, and reward magnitude. For all our analyses, we used the hex-level summary dataset (hexLevelDf, 298,918 observations across 10 rats).

In the second dataset, used to characterise tonic DA signals in the absence of movement, Floeder et al.^28^ recorded NAc dopamine using dLight fiber photometry in head-fixed mice undergoing Pavlovian conditioning. Mice (n = 18) completed 16–24 sessions each, alternating between short-ITI (∼15 s median inter-trial interval) and long-ITI (∼45 s) conditions across days. On each trial, an auditory cue was followed by liquid reward delivery after a fixed delay. The dataset provided continuous 470 nm and 405 nm (isosbestic control) photometry signals, alongside timestamped event codes for cue onsets, reward deliveries, and lick events.

### Analysis

All data were analysed using the Julia programming language^29^ (v1.12). Anthropic Claude Code running Opus 4.6 was used as a coding and writing assistant. Numerical integration was performed using the software package DifferentialEquations.jl (Tsit5 solver)^30^, and figures were produced with CairoMakie.jl^31^. Significance threshold for all tests was set at p<0.05.

#### Krausz (2023) dataset (used in figures. 2, 3, and 5)

We included trials that were part of an approach epoch. Observations where the rat was not in transit, was at the reward port itself, had already received reward, or had missing values were excluded. This yielded 264,042 included observations across 10 rats. For analyses requiring separate moving and stationary estimates (Fig. 2G–H, Fig. 3), we excluded observations with intermediate velocities, retaining only moving (velocity ≥ 15 cm/s) and stationary (velocity < 5 cm/s) epochs.

DA was averaged first within each rat at each hex position, then across rats (mean ± SEM). For velocity and reward splits, we performed within-rat quartile splits: each rat’s observations were divided into fast/slow (top/bottom quartile of velocity) or high/low reward (destination reward ≥ 80 vs ≤ 20 µL). For learning effects, sessions were split into early (first quartile) and late (last quartile) within each rat. For the hex-ranked analysis (Fig. 2G), we ranked the 49 maze positions by their mean moving-only DA as a proxy for spatial value, avoiding circularity from including stationary observations in the ranking. Stationary DA was tested against zero using one-sample t-tests across rats. Moving vs stationary comparisons used paired t-tests (per-rat mean). Within-position DA–velocity correlations were computed as Pearson’s r at each hex position.

For the value function recovery (Fig. 3), we exploited an algebraic identity of the continuous-time TD RPE. When moving, the predicted signal is 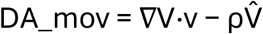; when stationary, v = 0, so 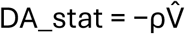. Subtracting the two eliminates the tonic term exactly, yielding DA_mov − DA_stat = ∇V·v. We estimated the value gradient at each hex position as ∇V = (DA_mov − DA_stat) / v, where DA_mov and DA_stat are the mean DA for moving and stationary observations at that position, and v is the mean moving velocity. We then integrated ∇V numerically along the approach axis using the Runge– Kutta solver (Tsit5) to recover V(x) up to an additive constant. To characterise the shape of the recovered value function, we fit four candidate profiles to each rat — exponential, hyperbolic, power-law, and Gaussian — each with three free parameters, and selected the best-fitting shape using BIC.

#### Floeder (2025) dataset (Fig. 4)

We extracted photometry signals, event timestamps, and lick events from NWB files (n=18 mice). The dLight signal (470 nm) was corrected for motion artifacts by regressing out the isosbestic control signal (405 nm) and computing ΔF/F. ITI condition (short vs long) was assigned per session based on the median inter-cue interval (< 25 s = short, ≥ 25 s = long).

To measure the tonic DA dip during the ITI (Fig. 4), we binned the ΔF/F signal in 1s windows locked to reward delivery, truncating each trial’s trace 1s before the next cue onset to prevent cross-trial contamination. Bins with fewer than 20 observations were excluded. For each subject, we computed the mean ΔF/F over the second half of the ITI (from the midpoint to 1s before the next cue) in long-ITI trials with ITI duration ≥ 30 s, yielding a per-session tonic DA measure. Anticipatory lick rate was computed as the number of lick onsets in the 5–8 s window after cue onset (before typical reward delivery), divided by 3s. To test whether the tonic dip deepens and vigour increases with learning, we regressed these measures against session number using linear mixed-effects models with random intercepts and slopes per subject. Short vs long ITI anticipatory lick rates were compared using paired t-tests across subjects.

#### Model Comparison (Fig 5)

*Model comparison (Fig. 5)*. We compared six functional forms derived from competing accounts of dopamine during navigation. Each model specifies a predicted DA as a function of distance to goal (d) and velocity (v):

- CT TD (continuous temporal difference exponential): *Ae*^− *kd*^*v* + *Be*^− *kd*^ + *c*
- CT TD (Gaussian): 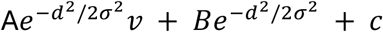
- dV/dt (exponential): *Ae*^− *kd*^*v* + *c*
- dV/dt (Gaussian): 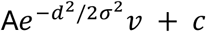
- Value: *Ae*^− *kd*^ + *c*
- Velocity: *Av* + *c*

The CT TD models had 4 free parameters, the dV/dt and Value models had 3, and the Velocity model had 2. Models were fit to each rat’s approach observations separately using Nelder–Mead optimisation with multiple starting points (4 initial parameter vectors plus 10 random perturbations each; 200,000 iterations per start). The best fit (lowest sum of squared errors) across all starts was retained. Model comparison used BIC. We report ΔBIC relative to the best-fitting model, summed across rats. Further detailed methods are reported in the supplementary information.

## ACKNOWLEDGEMENTS

We thank Thomas Akam, Luke Priestley, Sanjay Manohar for their helpful comments on our manuscript.

## AUTHOR CONTRIBUTIONS

S.G. derived the framework, analysed data, and wrote the manuscript. LM supervised the research, and wrote the manuscript.

## DATA AVAILABILITY

The Krausz et. al (2023) dataset is publicly available at 10.17632/m59zdjpm9h.1. The Floeder et. al (2025) dataset is publicly available at 10.48324/dandi.001552/0.250828.1728

## CODE AVAILABILITY

Code used to generate the figures is available at https://osf.io/g3e2d/

